# Transcriptional responses of *Biomphalaria pfeifferi* and *Schistosoma mansoni* following exposure to niclosamide, with evidence for a synergistic effect on snails following exposure to both stressors

**DOI:** 10.1101/446310

**Authors:** Sarah K. Buddenborg, Bishoy Kamel, Si-Ming Zhang, Gerald M. Mkoji, Eric S. Loker

## Abstract

**Background:** Schistosomiasis is one of the world’s most common NTDs. Successful control operations often target snail vectors with the molluscicide niclosamide. Little is known about how niclosamide affects snails, including for *Biomphalaria pfeifferi*, the most important vector for *Schistosoma mansoni* in Africa. We used Illumina technology to explore how field-derived *B. pfeifferi*, either uninfected or harboring cercariae–producing *S. mansoni* sporocysts, respond to a sublethal exposure of niclosamide. This study afforded the opportunity to determine if snails respond differently to biotic or abiotic stressors, and if they reserve unique responses for when presented with both stressors in combination. We also examined how sporocysts respond when their snail host is exposed to niclosamide.

**Principal Findings:** Cercariae-producing sporocysts within snails exposed to niclosamide express ~68% of the genes in the *S. mansoni* genome, as compared to 66% expressed by intramolluscan stages of *S. mansoni* in snails not exposed to niclosamide. Niclosamide does not disable sporocysts nor does it seem to provoke from them distinctive responses associated with detoxifying a xenobiotic. For *B. pfeifferi*, niclosamide treatment alone increases expression of several features not up-regulated in infected snails including particular cytochrome p450s and heat shock proteins, glutathione-S-transferases, antimicrobial factors like LBP/BPI and protease inhibitors, and also provokes strong down regulation of proteases. Exposure of infected snails to niclosamide resulted in numerous up-regulated responses associated with apoptosis along with down-regulated ribosomal and defense functions, indicative of a distinctive, compromised state not achieved with either stimulus alone.

**Conclusions/Significance:** This study helps define the transcriptomic responses of an important and under-studied schistosome vector to *S. mansoni* sporocysts, to niclosamide, and to both in combination. It suggests the response of *S. mansoni* sporocysts to niclosamide is minimal and not reflective of a distinct repertoire of genes to handle xenobiotics while in the snail host. It also offers new insights for how niclosamide affects snails.

**Author’S Summary:** Schistosomaisis control programs often employ the use of chemical molluscicides, such as niclosamide, to control the obligatory intermediate snail hosts. Despite its widespread use, we know little about how niclosamide affects snails like *Biomphalaria pfeifferi*, the most important vector *Schistosoma mansoni* in Africa. By sequencing the transcriptomes of uninfected and *S. mansoni*-infected *B. pfeifferi* exposed to niclosamide, we analyze the snail’s response to both biotic and abiotic stressors. We can also examine the response of *S. mansoni* to niclosamide exposure during intramolluscan development. *Biomphalaria pfeifferi* snails exposed only to niclosamide showed unique up-regulation of stress and defense-related transcripts not seen in snails infected with a biotic, like *S. mansoni* infection, alone. *Schistosoma mansoni*-infected *B. pfeifferi* exposed to niclosamide were clearly unable to regulate normal metabolic and detoxification processes. Cercariae-producing sporocysts within snails exposed to niclosamide are largely unaffected and continue to produce transcripts required for cercariae production.

## INTRODUCTION

Schistosomiasis control remains elusive in many of the world’s hyperendemic foci of infection in sub-Saharan Africa, jeopardizing the goals of diminishing schistosomiasis as a public health concern, or of eliminating transmission where possible by 2025 [1]. Several recent papers have called for the need to adopt more integrated control approaches instead of relying on chemotherapy alone to achieve eventual elimination [2–3], and there has been a resurgence in interest in methods to control the snails that vector human schistosomiasis [4–5]. Although the practical options available for use in snail control remain limited, molluscicides have been advocated because there are several recorded instances where their use has been associated with successful control [4,6].

Following the discovery of niclosamide’s molluscicidal properties in the 1950s, it has been incorporated into the commercial preparation known as Bayluscide [7] and is the only molluscicide approved for use in schistosomiasis control by the WHO Pesticide Evaluation Scheme (WHOPES). Use of niclosamide has enjoyed a modest resurgence and its focal application in snail control is advocated by WHO [8]. It has been used widely in Egypt and China as a mainstay for control operations, and it is used in both experimental [9–10] and in new control contexts, most notably recently as part of the *S. haematobium* elimination program in Zanzibar [11–12].

Although some work on the effects of molluscicides on oxygen consumption and carbohydrate metabolism of snails has been undertaken [13–14], there have been relatively few studies employing modern techniques to assess the impacts of molluscicide exposure on schistosome-transmitting snails. Zhao et al. [15], working with the amphibious snail *Oncomelania hupensis*, the intermediate host for *Schistosoma japonicum*, undertook an Illumina-based *de novo* transcriptome study to show this snail responded to two novel niclosamide-based molluscicides by up-regulating production of two cytochrome p450 (CYPs) genes, and one glutathione-S-transferase. Zhang et al. [16] examined the effects of three different sublethal concentrations of niclosamide (0.05, 0.10, and 0.15 mg/L for 24 hours) on the transcriptional activity of *Biomphalaria glabrata* as examined using an oligonucleotide microarray and noted up-regulation of several genes associated with biotransformation of xenobiotics (CYPs and glutathione-S-transferase), drug transporters, heat shock proteins (HSP 20, 40 and 70 families) and vesicle trafficking. Down-regulated hemoglobin production was also noted. Niclosamide is able to kill schistosome miracidia and cercariae [17–18] and field experiments in China have shown that niclosamide is effective at reducing the number of viable *S. japonicum* cercariae in streams and downstream infection of sentinel mice [19].

With respect to the effects of niclosamide on schistosome-infected snails, or on the schistosome sporocysts within them, there has been remarkably little study. Sturrock [20] investigated the effects of sublethal concentrations of niclosamide on infections of *S. mansoni* on *Biomphalaria sudanica tanganyicensis* and noted that: 1) snails exposed to molluscicide that survived were still susceptible to infection; 2) snails with prepatent infections were not initially more susceptible to molluscicide but had slightly delayed rate of parasite development and production of cercariae and did eventually exhibit higher mortality as they entered patency; and 3) survivorship of snails exposed during the patent period was less, although it takes some time for the effect to occur. Sturrock [20] commented that the combined stress of producing cercariae and exposure to molluscicide likely contributed to the higher mortality rate in patent snails. He also noted that doses sufficiently high to kill schistosome sporocysts in snails were probably above the lethal doses needed to kill the snails themselves.

In this study, building on the microarray results of Zhang et al. [16] with *B. glabrata*, we sought to obtain a more in-depth view of the transcriptome of molluscicide-exposed snails by using the Illumina platform to examine the responses of *Biomphalaria pfeifferi* to a sublethal dose (0.15 mg/L) of niclosamide. *Biomphalaria pfeifferi* is widely distributed in streams, ponds and impoundments in Africa and is probably responsible for transmitting more cases of *Schistosoma mansoni* than any other *Biomphalaria* species [21–22]. In addition, we examined the transcriptional responses to the same dose of molluscicide of *B. pfeifferi* harboring cercariae-producing *S. mansoni* infections. We were able to compare the responses of the above snails to both uninfected and infected *B. pfeifferi* not exposed to molluscicides (see companion studies [23,24]). For both the previous and present studies, we chose to examine the responses of snails recently removed from field habitats and therefore considered to be more representative of what might be expected of snails comprising natural populations actually exposed to molluscicides. The approach taken enables us to ascertain if and how the transcriptional responses of snails already coping with a massive *S. mansoni* infection can be further altered by simultaneous exposure to a toxic xenobiotic. For example, might snail genes up-regulated following exposure to *S. mansoni* trend towards down-regulation if the snail is exposed to niclosamide and required to produce increased quantities of molecules involved in detoxification?

With respect to the sporocysts of *S. mansoni* residing in snails exposed to niclosamide, do they exhibit any tendency to express genes that are not normally expressed during intramolluscan development, and if so, do the ensuing proteins favor survival of the sporocysts or of the stressed snail in which the sporocysts reside? Three possible scenarios for *S. mansoni* transcriptional response to molluscicide exposure can be considered: 1) We see an overall down-regulation of *S. mansoni* transcripts indicating suspension of activity; 2) Cercariae-producing *S. mansoni* sporocysts express unique features that are shut off in response to molluscicide exposure; and 3) Shedding *S. mansoni* stages exposed to molluscicide show unique transcriptional responses suggestive of a hitherto unseen ability to protect the host-parasite unit in which they reside from a xenobiotic.

## METHODS

*Biomphalaria pfeifferi* used in Illumina sequencing were collected from Kasabong stream in Asembo Village, Nyanza Province, western Kenya (34.42037°E, 0.15869°S) and transferred to our field lab at The Centre for Global Health Research (CGHR) at Kisian, western Kenya. Snails sized 6-9mm in shell diameter were placed under natural light to check for shedding of digenetic trematode cercariae [25]. Snails shedding only *S. mansoni* cercariae and non-shedding controls were used in this study. More details of collections and processing can be found in Buddenborg et al. [23–24].

The molluscicide niclosamide dissolved in dimethyl sulfoxide (DMSO) was purchased from Sigma. Uninfected and S. *mansoni-infected B. pfeifferi* were exposed to a concentration of 0.15 mg/L niclosamide with final DMSO concentrations at 1/1000 (v/v) for 24 hours at 26-28°C with aeration [16]. Previous 24 h exposure of *B. glabrata* to varying doses of niclosamide (0.05mg/L, 0.10mg/L, and 0.15mg/L) found that the 0.15mg/L dose produced the most robust transcriptional response, as assessed by microarray analysis [16]. All snails exposed to 0.15mg/L niclosamide were alive and responding after the 24 hours dosage period. Therefore, a 0.15mg/L dose was also selected for this study as the sublethal dose administered to *B. pfeifferi.*

RNA extraction, library preparation, sequencing procedures, and sequencing summaries can be found in Buddenborg et al. [23,24]. Illumina RNA sequencing reads underwent extensive processing in order to separate host, parasite, and potential symbiont reads. *Biomphalaria pfeifferi* read quantification and differential expression analyses were performed using RSEM (RNA-Seq by expectation maximization) [26] and EBSeq [27]. Normalized *Schistosoma mansoni* read counts acquired from RSEM were normalized using DESeq’s median of ratio method [28].

## RESULTS AND DISCUSSION

### Overall *B. pfeifferi* and *S. mansoni* transcriptomic responses to molluscicide exposure

Relative to uninfected and untreated control *B. pfeifferi*, the overall differential gene expression responses were measured for snails i) with shedding *S. mansoni* infections only, ii) exposed for 24 h to a sublethal dose of niclosamide only, or iii) harboring shedding *S. mansoni* infections *and* exposed to niclosamide (Fig 1A). The responses of shedding snails relative to unexposed controls have been discussed extensively by Buddenborg et al. (2017). With respect to molluscicide exposure, this is the first Illumina-based view of the transcriptomics response for any species of planorbid snail, and supplements and extends the view provided by the microarray study for uninfected *B. glabrata* of Zhang et al. [16]. Zhao et al. [15] undertook an Illumina-based study of the molluscicide-induced transcriptome of *Oncomelania hupensis*, the pomatiopsid snail host of *S. japonicum*. The response of *B. pfeifferi* to simultaneous exposure to schistosome infection and niclosamide exposure is the first glimpse we have for how snails respond transcriptionally to simultaneous exposure to these two relevant stressors.

**Fig 1.**
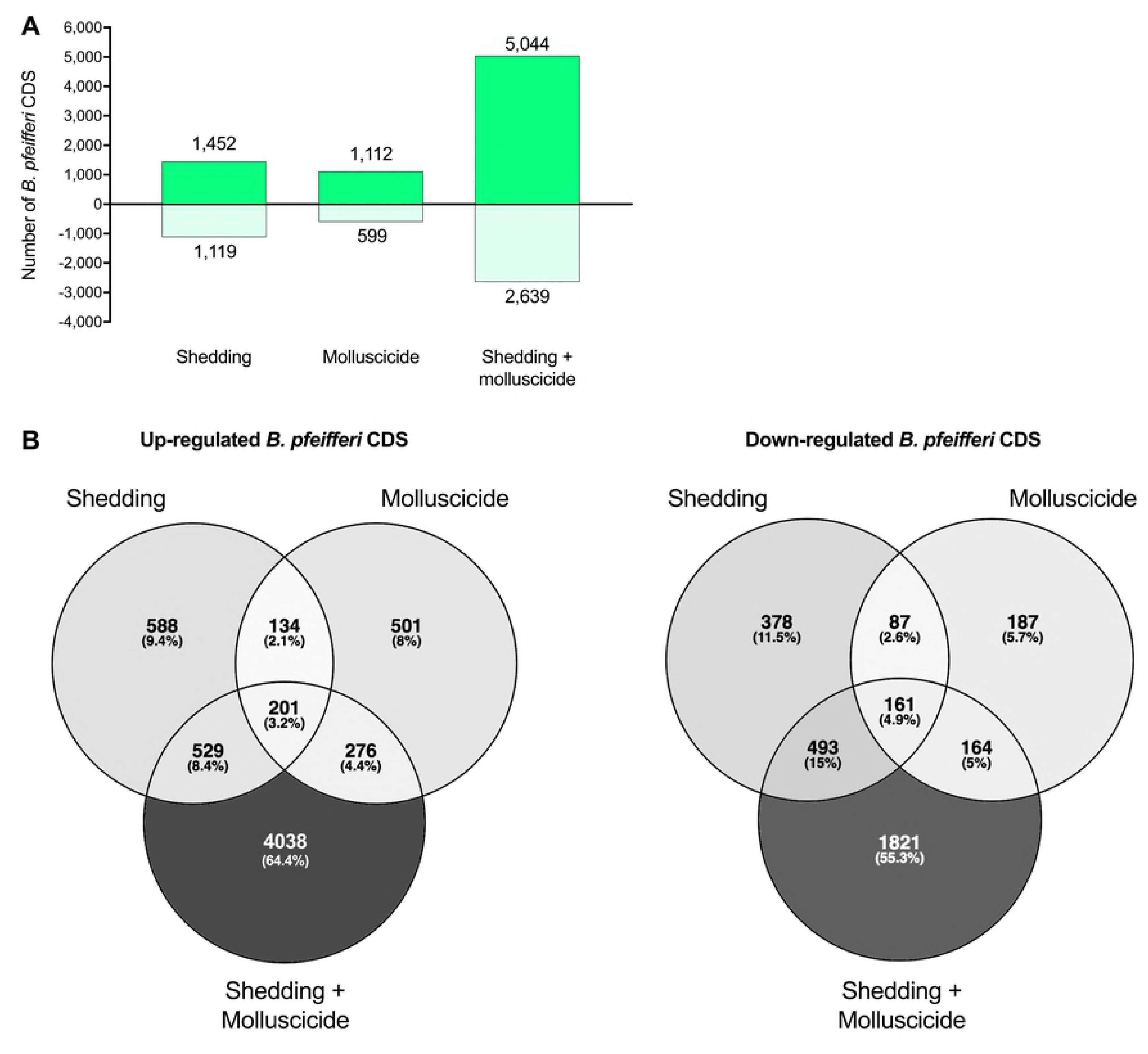
(A) Number of *B. pfeifferi* CDS up- and down-regulated in shedding and shedding plus molluscicide-exposed snails when compared to uninfected snails. (B) Venn diagrams showing shared and unique *B. pfeifferi* CDS between differentially expressed groups.

For each of the three treatments noted, the number of upregulated snail features exceeded the number of down-regulated features. For both up- and down-regulated features, it was remarkable that over half of the transcripts proved to be distinctively represented in the combined treatment group (Fig 1B). Over 4,000 genes were distinctively up-regulated in the snails receiving the combination of stressors. This was the largest number found in any single group of either venn diagram. It was surprising to us that larger numbers of genes were not found in the cells of either venn diagram that represented two or all three of the treatments. It was also evident that although the response of niclosamide-exposed snails had features in common to those evoked by *S. mansoni* exposure, many genes were also uniquely differentially expressed by exposure to just niclosamide. Further inspection of the pattern in expression levels exhibited by genes uniquely expressed in the combined treatment group revealed that in comparison to genes represented in other cells, they were modest in the degree of their differential expression. The specific nature of the genes responsive to either molluscicide alone, or to molluscicides and *S. mansoni* are discussed further below.

The transcriptomic responses of intramolluscan stages of *S. mansoni*, including those from snails actively shedding cercariae are described by Buddenborg et al. [24], and are supplemented here by responses of shedding snails exposed to niclosamide (Fig 2A). *S. mansoni* from shedding snails expressed 18,736 transcripts whereas *S. mansoni* from shedding samples exposed to sublethal niclosamide expressed 23,040 transcripts (Fig 2B), with the majority (80.6%) of these shared between the two groups. Most of the remaining transcripts were unique to *S. mansoni* from the niclosamide-exposed samples (19%), with only 0.4% expressed uniquely in the shedding snails. The additional genes expressed only in the presence of niclosamide raises the percentage of the *S. mansoni* genome of 66% shown by Buddenborg et al. [24] to be expressed in snails to 68%. *S. mansoni* exposed to niclosamide expressed more transcripts, but this response was variable among replicates and in general most (>90%) of these extra transcripts were expressed less than 2 log_2_ normalized counts when replicate counts were averaged (Fig 2C). Of the 80.6% of shared *S. mansoni* transcripts, there was little difference in overall expression levels for transcripts from samples with or without niclosamide exposure.

**Fig 2.**
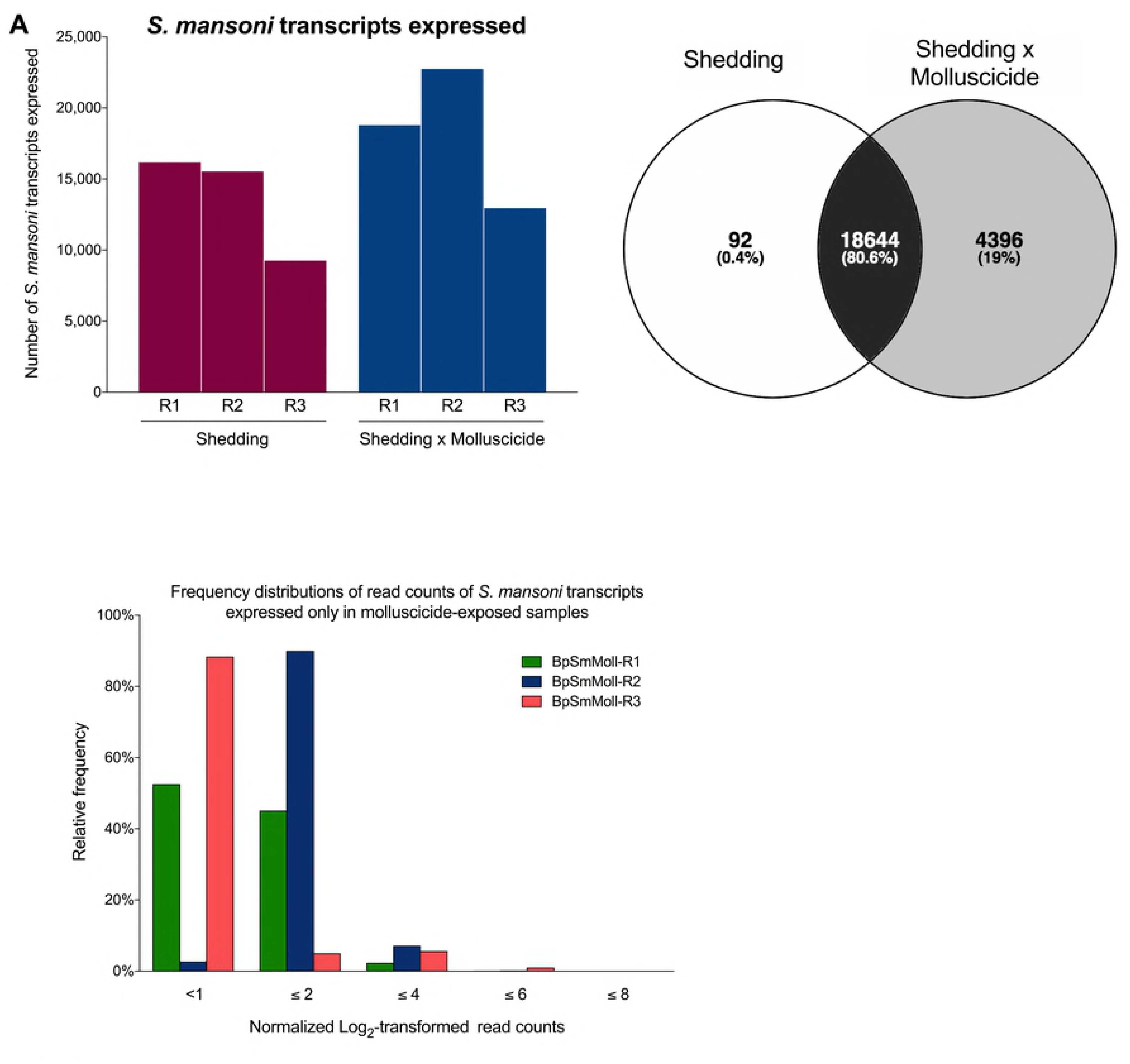
(A) *S. mansoni* transcripts expressed per replicate in shedding and shedding plus molluscicide exposed *S. mansoni*. (B) Venn diagram of shared and unique *S. mansoni* transcripts in tested groups. (C) Frequency distribution of log_2_-transformed normalized read counts of *S. mansoni* transcripts uniquely expressed in shedding plus molluscicide replicates.

### Specific responses of *S. mansoni* cercariae-producing sporocysts within *B. pfeifferi* exposed for 24 h exposure to sublethal niclosamide treatment

Sturrock [20] noted that the lethal dose of niclosamide for intramolluscan schistosomes must be higher than what is needed to kill the host snail. This is likely true since *S. mansoni* sporocysts are protected both by their syncytial tegument [29] and by being embedded in the host snail’s tissues. At least with respect to a 24 h exposure to a dose of niclosamide sublethal for *B. pfeifferi*, we observed that sporocysts within such snails produced more transcripts relative to sporocysts from untreated shedding snails, not fewer (Fig 2). If cercariae-producing sporocysts were strongly and directly affected by niclosamide, we would have expected to see extensive and broad down-regulation or absence of numerous transcripts such as those related to nutrient uptake across the tegument, of elastases indicating a decrease or pause in cercariae production, and of transcripts associated with germ ball development and proliferation. As it was, only 0.4% of transcripts were missing as compared to untreated sporocysts.

Defense and stress responses were sustained in niclosamide-treated sporocysts relative to untreated sporocysts (Fig 3). Peroxiredoxins like glutathione peroxidase and thioredoxin peroxidase, which may be responsible for elimination of potentially lethal hydrogen peroxide produced by the snail [30–32], were stably maintained. Stable and consistent expression of *S. mansoni* planarian-like bacterial defense homologs, heat shock proteins and SODs are all indicative that protective responses were maintained following exposure to niclosamide. Interestingly, there was no conspicuous evidence of *greater* representation of these transcripts in sporocysts from snails treated with niclosamide. Particularly noteworthy is the lack of an obvious response of the single *S. mansoni* cytochrome p450 gene to niclosamide presence. As noted by Ziniel et al. [33], the exact function of this gene product in *S. mansoni* is not known, but our results suggest it is not provoked by a xenobiotic like niclosamide. As noted by Ziniel et al. [33] and by us previously (Buddenborg et al. [24], parasitic helminths in general lack extensive cytochrome p450 repertoires, quite unlike their hosts. In contrast, *B. pfeifferi* does indeed deploy cytochrome p450 responses upon exposure to molluscicides (see below). Another group of *S. mansoni* transcripts of potential relevance to their response to niclosamide are drug efflux transporters like ABC transporters which are known to be up-regulated in adult schistosomes exposed to praziquantel [34–35]. We noted several efflux transporters were expressed in cercariae-producing sporocysts of *S. mansoni* [24] but did not observe any conspicuous change in their expression pattern following exposure to niclosamide (Fig 4). A study on the giant liver fluke *Fasciola gigantica* exposed to rhodamine-labeled niclosamide did not reveal substantial changes in ABC transporter activity although the authors did not rule out the potential involvement in these proteins in drug resistance and detoxification [36].

**Fig 3.**
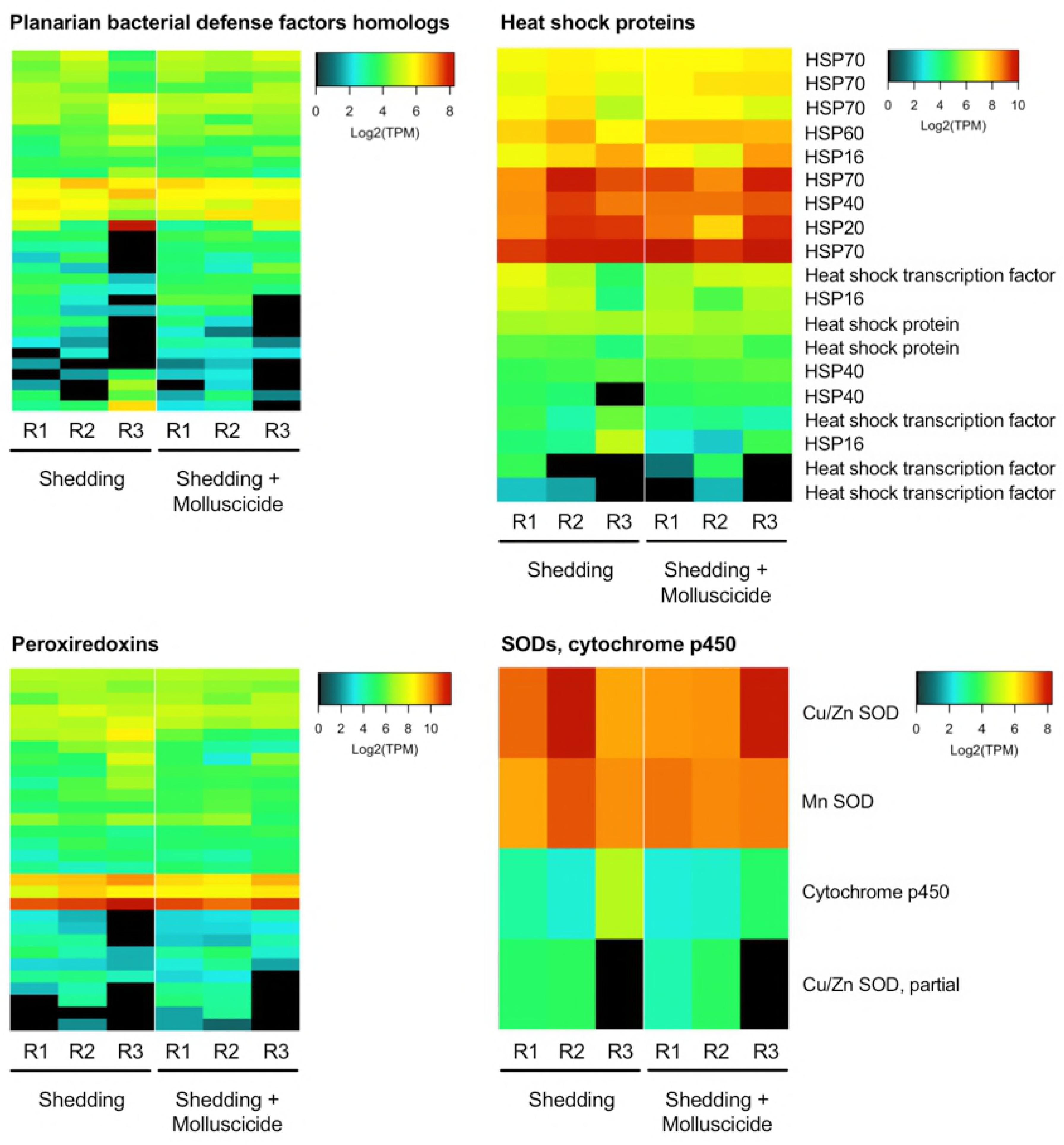
Stress and defense transcripts expressed by *S. mansoni*

**Fig 4.**
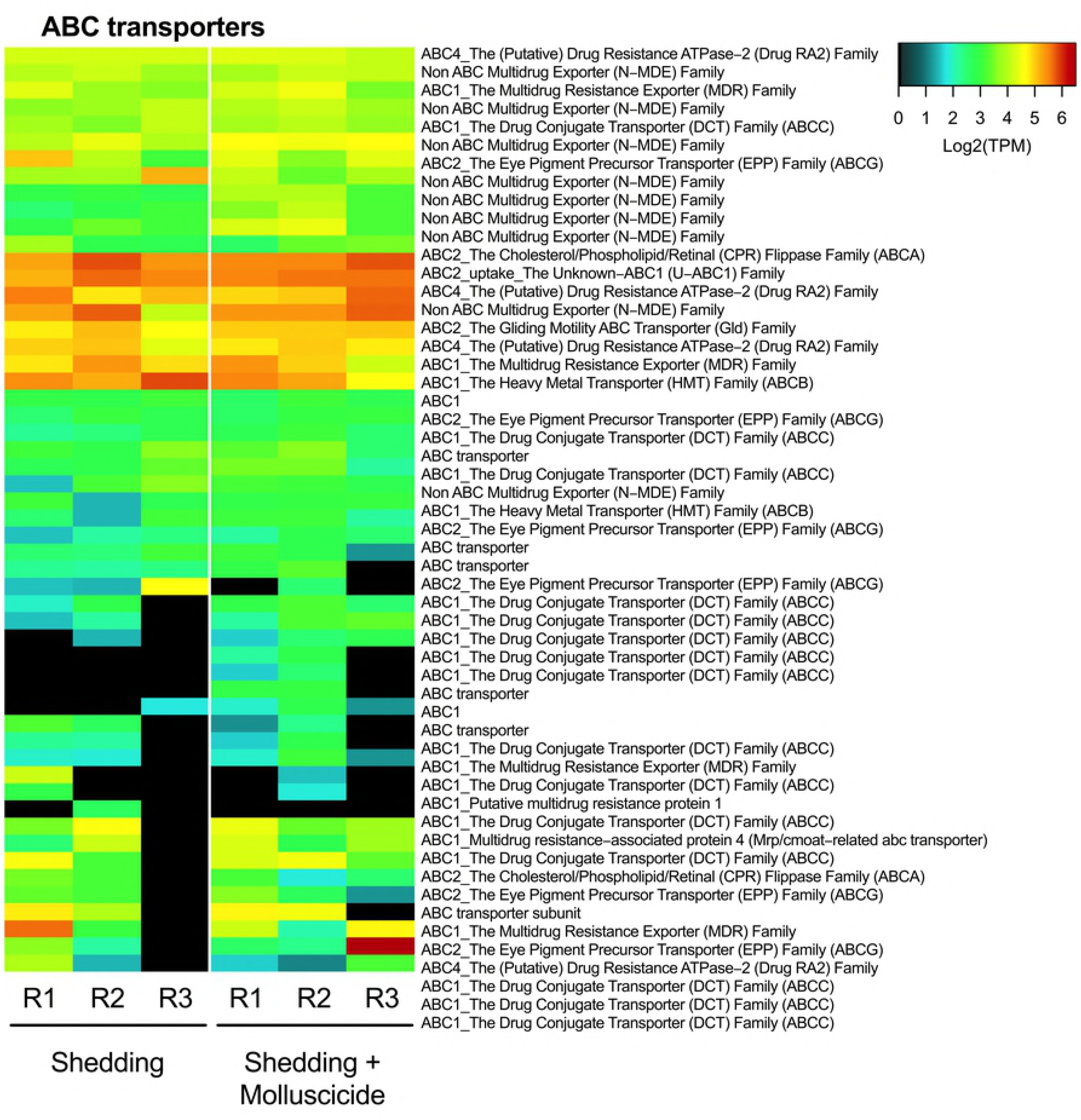
ABC transporters expressed by *S. mansoni* following exposure to 0.15mg/L niclosamide

Inspection of the transcripts produced uniquely by niclosamide-exposed sporocysts does not reveal any candidates that would seem to favor resilience to niclosamide. This coupled with the stable expression of known defense or stress response genes noted above leads us to a conclusion that sporocysts have little if any ability to mount protective responses to niclosamide and certainly do not seem to provide anything that would favor enhanced survival of their host snail in the presence of a chemical that is clearly lethal for the host. It is possible that the parasite is relying on host xenobiotic capabilities to respond to niclosamide.

*S. mansoni* sporocysts, and the cercariae developing within them, express a diverse array of proteases, including elastases and leishmanolysins [24,37], with likely functions in disabling snail defenses, dissolution of snail tissues to provide living space, facilitating intra-snail migration of sporocysts and for packaging in cercariae which use them both for exiting the snail host and entering the mammalian definitive host. Protease inhibitors are also produced and likely counteract proteases that the snail expresses late in infection [24,38]. The overall patterns of expression of proteases or protease-inhibitors does not differ substantially between sporocysts in untreated and niclosamide-treated snails (Fig 5).

**Fig 5.**
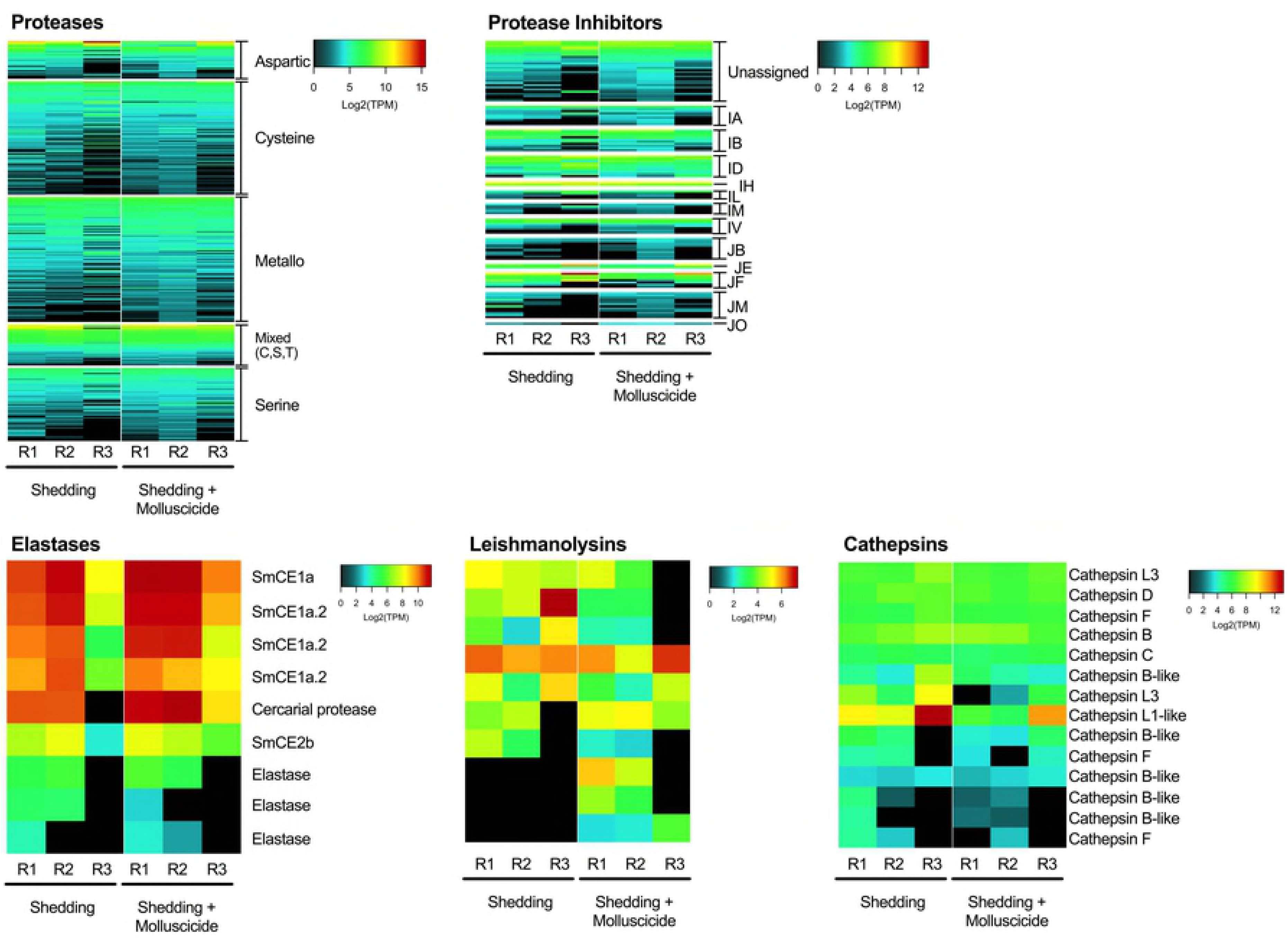
*Schistosoma mansoni* proteases and protease inhibitor transcripts expressed in shedding and shedding plus molluscicide groups.

When specific proteases like elastases, leishmanolysins, and cathepsins are examined, some changes in expression were noted. Niclosamide-exposed sporocysts exhibited modest increases in expression of cercarial elastases SmCE1a, SmCE1a.2, cercarial protease, and SmCE2b (Fig 5). The SmCE isoforms represent an expanded family of elastases unique to some *Schistosoma* species including *S. mansoni* [37]. Transcripts for leishmanolysins which are metalloproteases also known as invadolysins [39], also exhibited a modest increase in representation in niclosamide-exposed sporocysts as compared to untreated sporocysts [24].

At least some *S. mansoni* glucose, amino acid, and nucleoside transporter transcripts showed modestly increased representation in molluscicide-exposed sporocysts relative to untreated controls (Fig 6), as did some of the markers for germinal cell proliferation [40] such as fibroblast growth factor receptor 2, *vasa*, and *nanos-2* noted in Buddenborg et al. [24] (Fig 7). Very modest changes in expression were also noted (Fig 8) in neuropeptide hormones or markers of neural development important in flatworm locomotion, feeding, host location, regeneration, and development [41,42]. Shedding *S. mansoni* stages exposed to niclosamide had higher transcript levels for cell polarity protein, neuronal differentiation, notch, SOX transcription factor, and septate junction protein (Fig 7) and although modest, these may have important downstream effects on germinal cell proliferation or neurogenesis [43–45].

**Fig 6.**
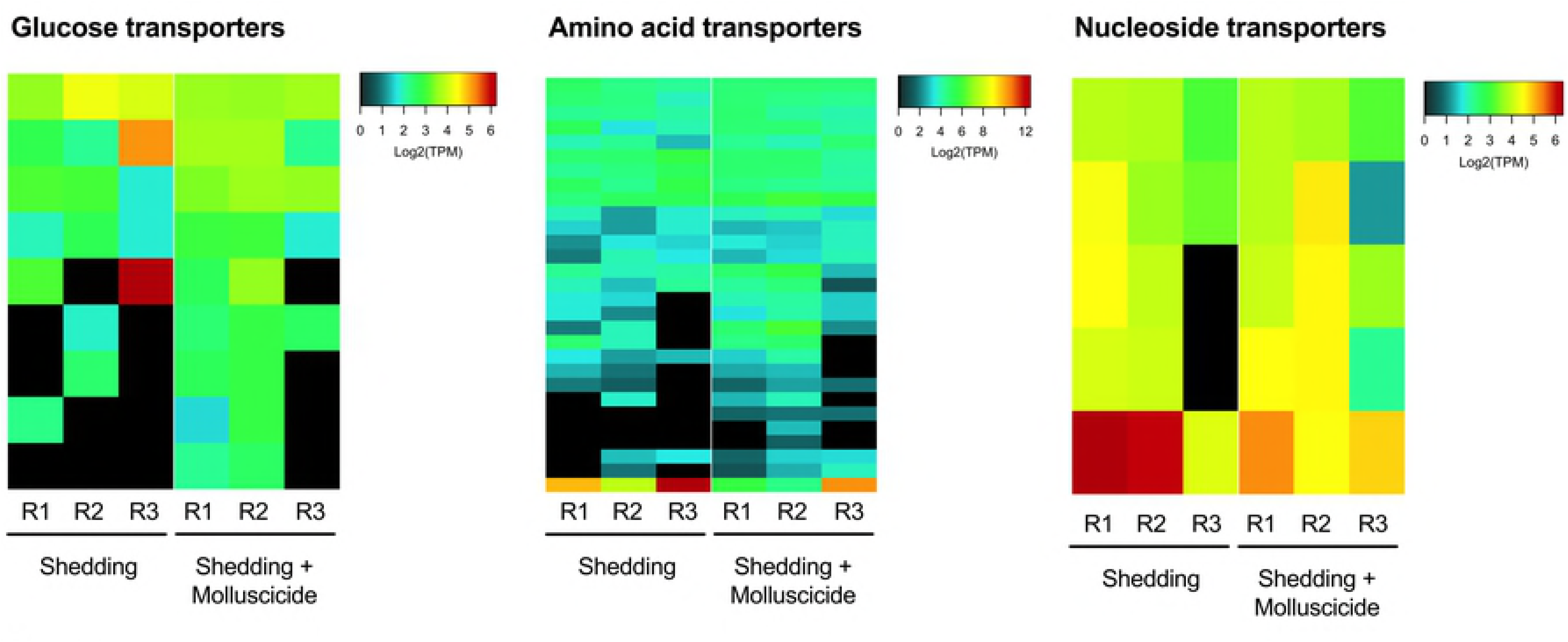
Nutrient transporter expression in shedding and shedding plus molluscicide-exposed *S. mansoni*.

**Fig 7.**
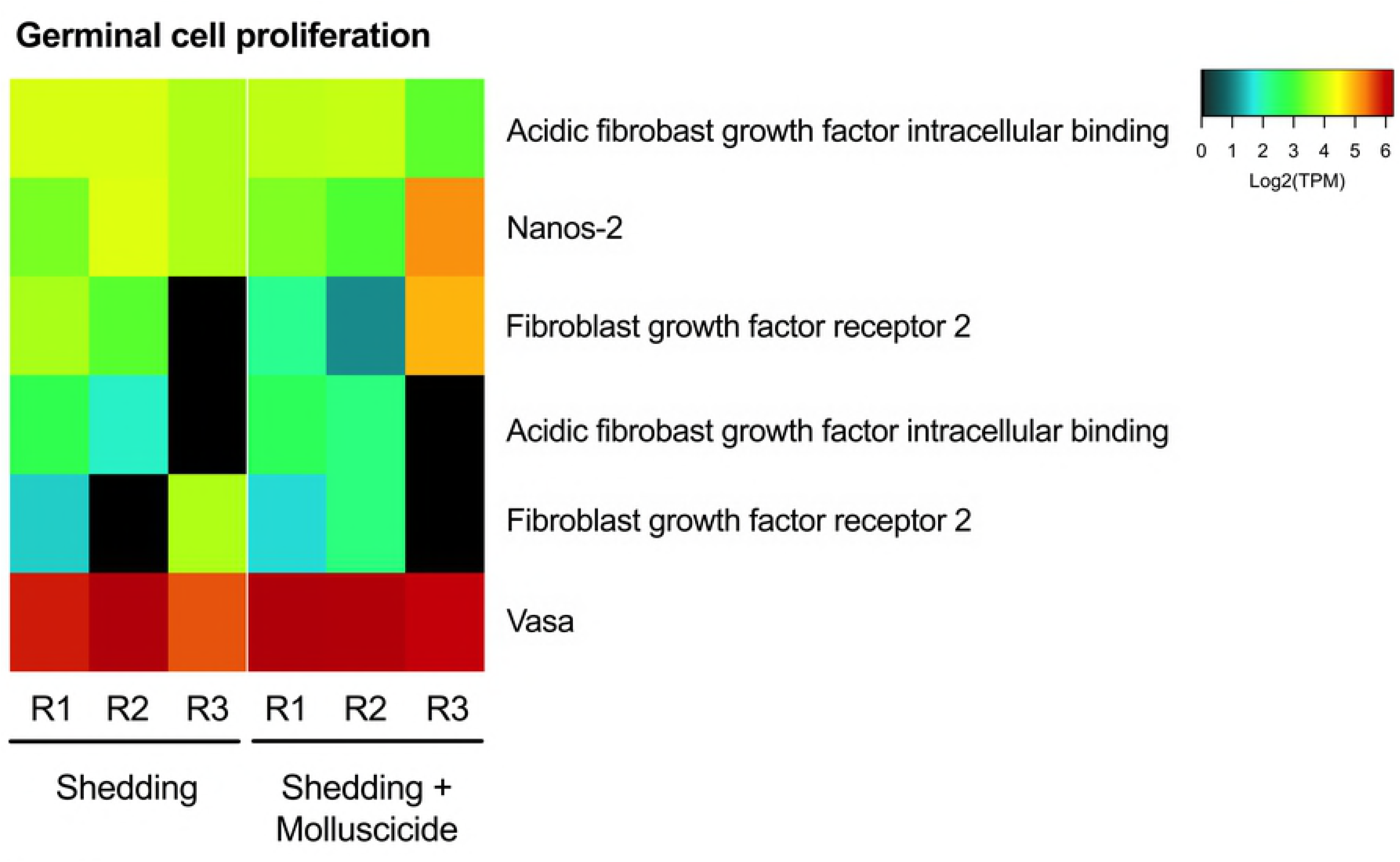
Expression of known germinal cell proliferation transcripts.

**Fig 8.**
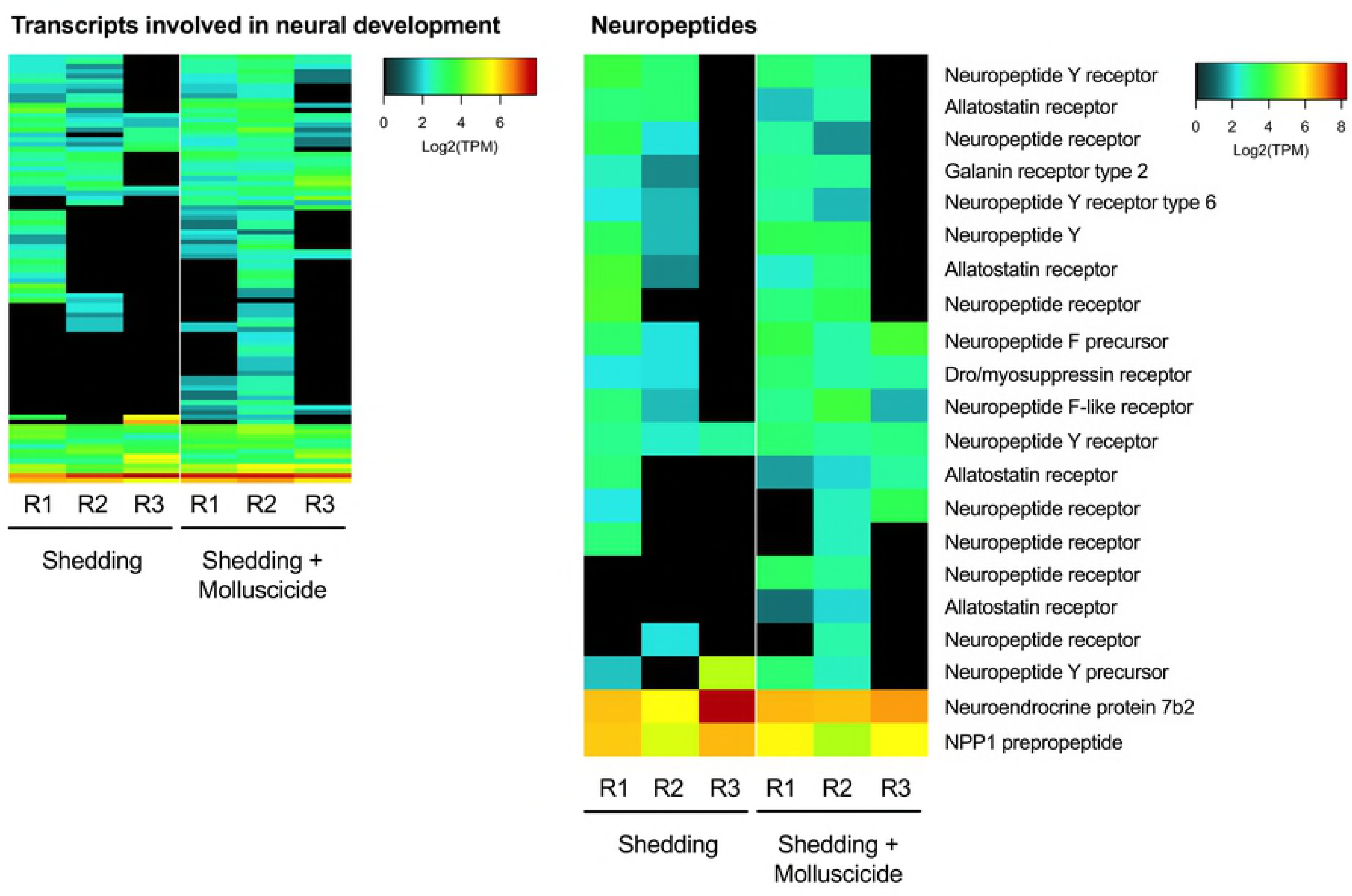
*Schistosoma mansoni* transcripts involved in neural development or encoding neuropeptides

The modest increases in proteases, transporters, germinal cell proliferation factors and neuropeptide or neural development markers in niclosamide-exposed *in vivo* sporocysts all serve to further highlight the fact that the 24 h niclosamide exposure we used was certainly not lethal to the sporocysts nor did it seem to significantly curtail their transcriptional production or to invoke transcripts associated either with enhanced efflux or processing of niclosamide or with apoptosis or autolysis of sporocysts. Of course, more extensive exposure of *B. pfeifferi* to niclosamide with attendant loss of the integrity of the snail metabolome would inevitably result in death of *S. mansoni* sporocysts as well.

### Shared response of two *Biomphalaria* species to a sublethal dose of niclosamide

Of the 30,647 probe features on the *B. glabrata* microarray used by Zhang et al. [16], 16,713 (55%) were homologous to a *B. pfeifferi* transcript (Blastn E-value <1e-10, percent identity >75%). Microarray features with homologs to *B. pfeifferi* transcripts and that were differentially expressed in both Zhang et al. [16] and the present study are shown in Table 1. These features represent a conservative view of genes characteristic of *Biomphalaria’s* response to sublethal niclosamide exposure. The entire differential expression analysis of *B. pfeifferi’s* response to niclosamide showed 895 transcripts up-regulated and 604 down-regulated when compared to uninfected control *B. pfeifferi.*

**Table 1.** All features shared between *B. glabrata* [16] and *B. pfeifferi* that were significantly differentially expressed after exposed to 0.15mg/L niclosamide.

|  | *B. pfeifferi*<br>Illumina transcript | Log<sub>2</sub><br>FC | *B. glabrata*<br>array feature | Log<sub>2</sub><br>FC |
| --- | --- | --- | --- | --- |
| ADP-ribosylation factor 3-like | evgTRINITY_DN92963_c1_g2_i1 | 5.10 | c13901 | 4.73 |
| ADP-ribosylation factor 3-like | evgTRINITY_DN92963_c1_g1_i1 | 4.45 | c13901 | 4.73 |
| Solute carrier family 28 member 3-like | evgTRINITY_DN88027_c1_g1_i4 | 6.78 | c27272 | 1.99 |
| Multidrug resistance 1-like | evgTRINITY_BU_DN81217_c7_g4_i1 | 2.72 | contig_14304 | 1.48 |
| Multidrug resistance 1-like | evgTRINITY_DN90366_c3_g1_i2 | 5.26 | contig_14304 | 1.48 |
| HSP 12 | evgIcl\|G0WVJSS02FHD9K | 2.29 | contig_7431 | 3.79 |
| HSP 12 | evgIcl\|G0WVJSS02JB97J | 1.98 | contig_7431 | 3.79 |
| HSP 70 | evgTRINITY_GG_25613_c6_g1_i1 | 1.09 | BGC03909 | 3.64 |
| Solute carrier family 28 member 3-like | evgTRINITY_DN88027_c1_g1_i3 | 3.79 | c27272 | 1.99 |
| Cytochrome p450 | evgTRINITY_BU_DN81631_c8_g1_i1 | 1.05 | c14547_rc | 3.10 |
| Cytochrome p450 | evgTRINITY_DN93193_c20_g1_i1 | 2.88 | c8814 | 2.88 |
| Baculoviral IAP repeat-containing 3-like | evgTRINITY_BU_DN78979_c0_g1_i2 | 1.69 | c17676_rc | 2.14 |
| Nuclear protein 1-like | evgIcl\|HJ4YRIA01D0DSV | 1.28 | contig_4627 | 2.20 |
| Nuclear protein 1-like | evgIcl\|HJ4YRIA02HBZUN | 1.01 | contig_4627 | 2.20 |
| Growth arrest and DNA damage-inducible alpha-like | evgIcl\|G0WVJSS02G7JUO | 1.85 | contig_8438 | 1.39 |
| Alpha-crystallin B chain | evgIcl\|HJ4YRIA01ERORD | 1.21 | contig_2362_rc | 1.79 |
| Sequestosome-1-like | evgTRINITY_DN29609_c0_g1_i1 | 1.00 | BGC02302 | 1.57 |
| Glycogen-binding subunit 76A-like | evgTRINITY_DN70212_c1_g1_i1 | 0.92 | c14016_rc | 1.09 |
| Methionine synthase reductase-like | evgTRINITY_DN77579_c0_g1_i1 | 0.71 | c41473 | 1.00 |
| Glutathione-independent glyoxalase hsp3103 | evgTRINITY_DN92822_c15_g1_i1 | -1.08 | contig_3480 | -1.10 |
| Thymidine kinase, cytosolic-like | evgTRINITY_DN90310_c10_g1_i1 | -1.29 | contig_10981 | -1.35 |
| Uncharacterized | evgTRINITY_DN89789_c4_g2_i1 | 2.39 | contig_12514_rc | 3.19 |
| Uncharacterized | evgIcl\|G0WVJSS01A5WAX | 3.89 | contig_6337_rc | 4.16 |
| Uncharacterized | evgIcl\|G0WVJSS01DEUAY | 1.60 | contig_3100 | 2.29 |
| Uncharacterized | evgTRINITY_DN88565_c20_g1_i1 | 2.00 | contig_3944_rc | 1.38 |
| Uncharacterized | evgTRINITY_DN22835_c0_g1_i1 | 1.33 | BGC02491 | 1.02 |
| Uncharacterized | evgIcl\|G0WVJSS01DKS66 | 1.36 | c43865_rc | 1.40 |
| Uncharacterized | evgIcl\|G0WVJSS02ITT0P | 0.82 | contig_7634_rc | 1.16 |
| Uncharacterized | evgTRINITY_GG_16388_c0_g2_i1 | 0.86 | c13164_rc | 1.09 |
| Uncharacterized | evgTRINITY_DN84827_c0_g2_i1 | -0.81 | c8798_rc | -1.13 |
| Uncharacterized | evgTRINITY_DN93461_c7_g1_i1 | -1.49 | c1870 | -1.80 |

As a lipophilic xenobiotic, niclosamide would likely be eliminated in animals by increasing its hydrophilicty (phase 1 reaction), conjugating the phase I product with a charged chemical group (phase 2 reaction), and then removing it with the aid of a transmembrane transporter (phase 3 reaction) [46]. A key enzyme superfamily of heme-thiolate proteins responsible for initial phase I detoxification are the cytochrome p450s (CYPs). CYPs are found in all kingdoms of life and most commonly perform monooxygenase reactions adding one oxygen atom to the xenobiotic with the other oxygen atom reduced to water [46]. Zhang et al. [16] found that 9 of the features that were up-regulated ≥ 2-fold change following exposure to 0.15mg/L of niclosamide were CYPs. The *B. glabrata* genome has about 99 genes encoding heme-thiolate detoxification enzymes with tissue-specific expression patterns suggesting that CYPs serve specific biological processes [47].

CYPs are also up-regulated in *B. pfeifferi* in response to niclosamide exposure, including two in common with *B. glabrata* (Table 1) and 8 more as noted in Fig 9A, underscoring the importance of CYP mixed function oxidases in the snail response to niclosamide. Of the CYPs up-regulated in both snail species, one is a homolog of Cp450 3A2-like found in mouse liver cell microsomes which is responsible for oxidizing steroids, fatty acids, and xenobiotics. The other shared CYP is CYP 3A41-like. It is also microsomal and studies of vertebrate homologs indicate that glucocorticoids may exert control of CYP3A41 gene expression [48]. Modest down-regulation of one CYP in *B. glabrata* (CYP II f2) was also observed [16] and we similarly noted down-regulation of a CYP (1-like isoform X1) in *B. pfeifferi.* This supports the suggestion by Zhang et al. [16] that different members of the CYPs repertoire are likely to have different functions in *Biomphalaria* snails in response to diverse stimuli, including biotic challenges like *S. mansoni* or abiotic challenges like molluscicides.

**Fig 9.**
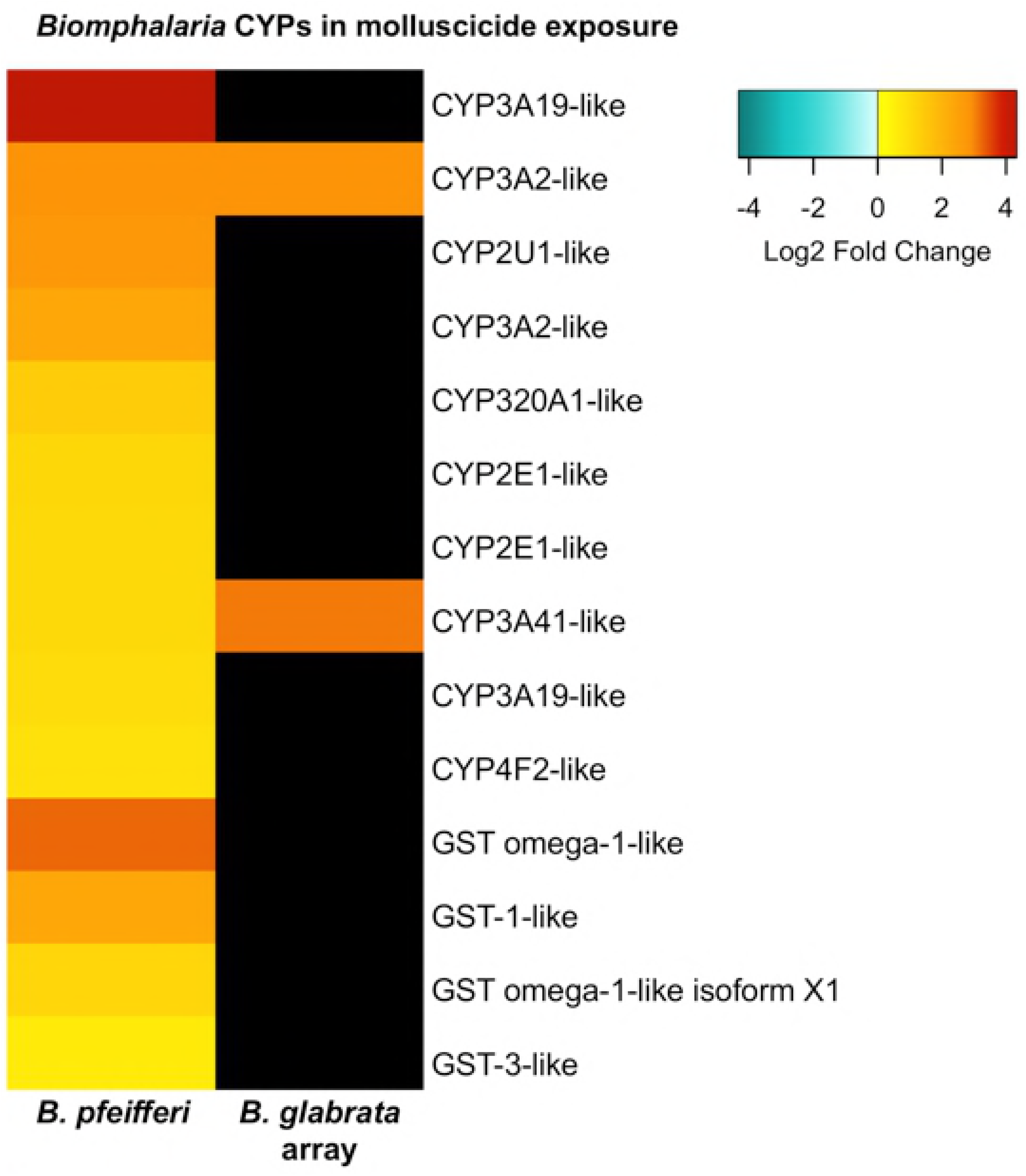
*Biomphalaria pfeifferi* CYP and GST transcripts up-regulated in response to niclosamide. Data for *B. glabrata* from Zhang et al. [16].

Phase 2 in the elimination of xenobiotics would likely include molecules like glutathione transferases that transfer charged chemical species like glutathione to the xenobiotic. Glutathione-S-transferase 7-like (GST) was up-regulated 5-fold following niclosamide exposure in *B. glabrata* [16]. In addition to CYPs, GST has also been shown to be up- regulated following niclosamide-based molluscicide exposure in *Oncomelania hupensis* [15]. GST was also represented in the *B. pfeifferi* Illumina DE transcripts with up-regulation of GST omega-1-like, and microsomal GST-1 and 3-like (Fig 9).

Transmembrane transporters complement the detoxification and conjugation reactions of phases 1 and 2 by eliminating the xenobiotic or toxin present in an organism [46]. ATP-binding cassette (ABC) transporters, particularly ABC efflux transporters, play an important role in eliminating toxic compounds from cells. For instance, ABCG2, a non-specific, multi-xenobiotic transporter is known to be expressed at high levels in the gills and hemocytes of *Mytilus edulis* [49]. One family of ABC efflux transporters, the multidrug resistance proteins (MRPs) act to eliminate drugs and toxic chemicals transporting anionic compounds detoxified in phases 1 and 2. One MRP-1 is expressed 2.8-fold higher than controls in *B. glabrata* [16] and 10 MRP-1 transcripts were up-regulated in *B. pfeifferi* suggesting these transporters are removing toxic waste products produced directly by niclosamide or indirectly through cell death or tissue necrosis (Table 2).

**Table 2.** *Biomphalaria pfeifferi* multidrug resistant protein 1-like isoforms up-regulated after exposure to sublethal niclosamide.

| *B. pfeifferi* Transcript | Log<sub>2</sub> FC |
| --- | --- |
| evgTRINITY_GG_18090_c5_g2_i1 | 7.8 |
| evgTRINITY_BU_DN63065_c0_g1_i1 | 5.7 |
| evgTRINITY_BU_DN81217_c7_g4_i3 | 5.7 |
| evgTRINITY_DN90366_c3_g1_i2 | 5.3 |
| evgTRINITY_DN90366_c3_g1_i1 | 5.0 |
| evgTRINITY_BU_DN81217_c7_g4_i1 | 2.7 |
| evgTRINITY_DN1870_c0_g1_i1 | 2.2 |
| evgTRINITY_DN92909_c10_g1_i1 | 1.4 |
| evgTRINITY_DN87672_c0_g1_i1 | 1.3 |
| evgTRINITY_DN84897_c0_g1_i1 | 1.3 |

Heat shock proteins show increased expressed to a variety of stressors including elevated temperature, hypoxia, ischemia, heavy metals, radiation, calcium increase, glucose deprivation, various pollutants, drugs, and infections [50]. Up-regulation of HSPs has been associated with susceptibility of *B. glabrata* to *S. mansoni* [51–52]. HSPs have also been identified in other molluscs as indicative of environmental stress. The disk abalone *Haliotis discus discus* up-regulates HSP 20 when exposed to extreme temperatures, changing salinity, hssseavy metals, and microbial infection [53]. The marine bivalve, *Mytilus galloprovincialis* up-regulates HSPs 24.1, 70, 90, and sequestosome-1 following toxic metal exposure [54]. *Biomphalaria glabrata* mounts a multifaceted HSP response to niclosamide by up-regulating HSPs 12, 40, and 70 [16] and the selective autophagosome cargo protein sequestosome-1. We also saw up-regulation of these specific HSPs but the more comprehensive sequencing available from the Illumina study revealed mixed responses of isoforms of HSP 12.2 and down-regulation of HSP 30 (Fig 10).

**Fig 10.**
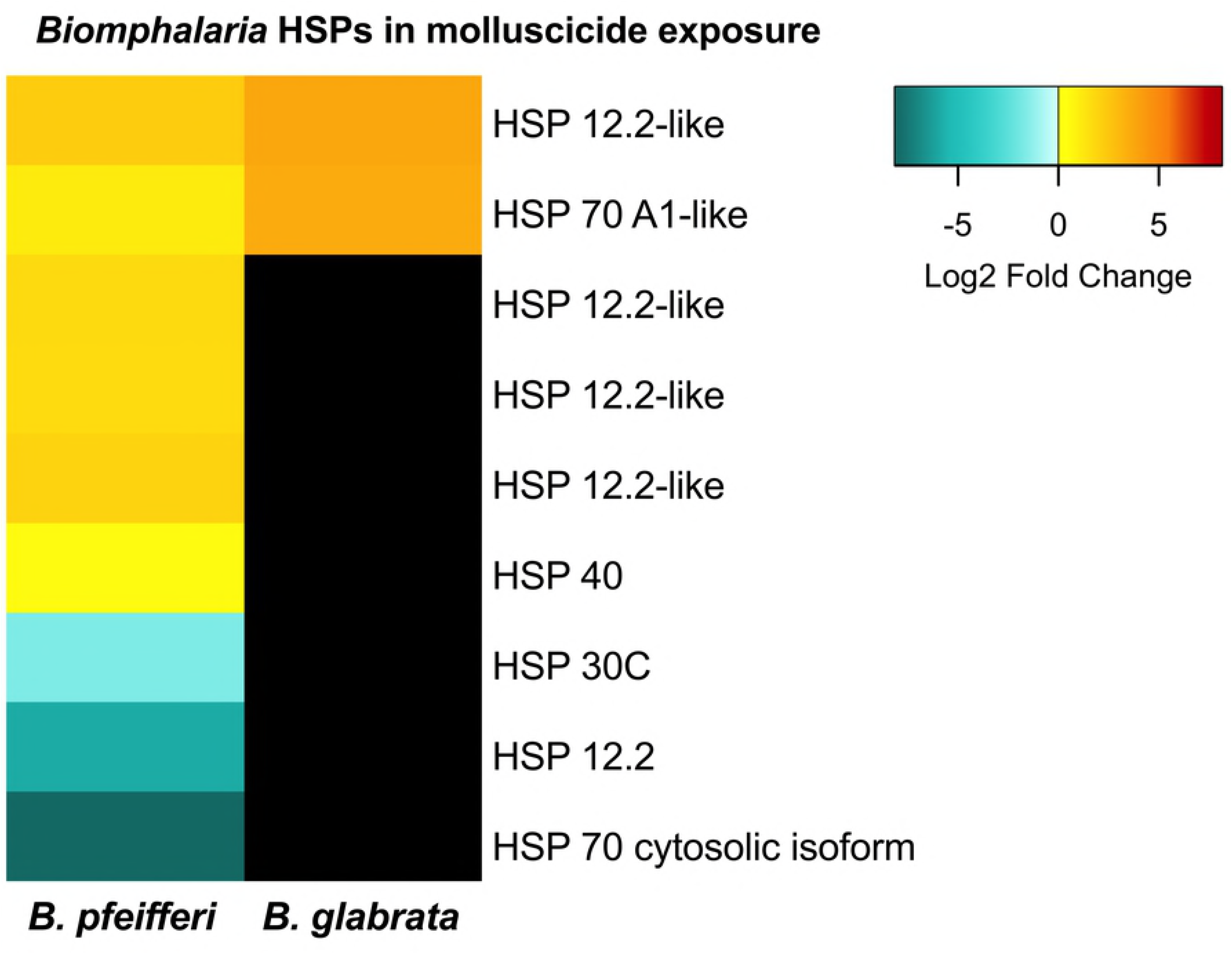
Expression of *B. pfeifferi* HSP transcripts and homologous *B. glabrata* HSP microarray probes in response to 0.15mg/L niclosamide. Data for *B. glabrata* from Zhang et al. [16].

In response to exposure to *S. mansoni* infection, *B. pfeifferi* shows a more complex transcriptional expression of HSPs, cytochrome p450s, and glutathione-S-transferases than it does to molluscicide with no general up- or down-regulation of any group of these transcripts [23]. Biotic stressors such as parasites with intimate and prolonged contact with host tissues may induce a more complex stress response with up- and down-regulation of various HSPs in comparison to a general up-regulated of CYPs, glutathione-S-transferases, small and large molecular weight HSPs, and sequestosome noted in the response of several molluscs to abiotic stressors.

### Additional responses of *Biomphalaria* to sublethal molluscicide exposure detected with Illumina RNA-Seq

Because of the unbiased sequencing available in Illumina RNA-Seq, we were able to acquire additional information on the transcriptomic responses of *B. pfeifferi* to niclosamide beyond that allowed by the *B. glabrata* microarray study. Transcripts involved in protection from oxidative damage, generalized pathogen defense and innate immunity, protease inhibitors and feeding behavior were all noted.

We observed high expression of several glutathione peroxidase transcripts, presumably associated with enhanced conversion of hydrogen peroxide to water. In cancerous colon cells, niclosamide increased cell death when used with a therapeutic drug through hydrogen peroxide production [55], therefore, it is not inconceivable that niclosamide in snails is directly or indirectly involved in increasing hydrogen peroxide levels although there is thus far no direct evidence for this in exposed snails. Glutathione peroxidase has been shown to increase the general tolerance of cells to oxidative stress resulting from exposure to xenobiotics [56].

Glutathione reductase, a critical oxidoreductase enzyme that catalyzes the reduction of glutathione disulfide to glutathione, surprisingly was down-regulated. As noted above, glutathione is a key ingredient needed in phase II conjugation mediated by the enzyme glutathione-S-transferase, which is up-regulated in *B. pfeifferi* following molluscicide exposure. An impaired ability to regenerate glutathione because of down-regulated glutathione reductase activity could then impair both the detoxification process and interfere with maintenance of redox balance by allowing hydrogen peroxide to accumulate.

One of the more striking responses of *B. pfeifferi* exposed to niclosamide was the high up-regulation of transcripts for several protease inhibitors including antitrypsin-like and serpins (serine protease inhibitors) and the down-regulation of metallo, cysteine, and serine proteases. In contrast, only one serine protease (chymotrypsin-like elastase family member 1) and a single aminopeptidase N-like transcript were up-regulated. Caspases are cysteine-dependent proteases that play essential roles in programmed cell death [57] and isoforms of caspase-2 and 3 were down-regulated in niclosamide-exposed *B. pfeifferi.* The down- regulation of protease activity may be part of a compensatory stress response made by the snail to minimize metabolic changes associated with niclosamide exposure that if left unchecked would lead to apoptosis and protein degradation.

Responses typically classified as innate immune responses because they occur following exposure to parasites like S. mansoni were also noted in *B. pfeifferi* exposed only to niclosamide. One such transcript was homologous to CD109 antigen-like, a thioester-containing protein, which is highly enriched in plasma from both resistant and susceptible strains of *B. glabrata* containing miracidia transforming into mother sporocysts [58]. We also noted up-regulation of a transcript identified as complement C1q-like protein that we have reported to be up-regulated in early S. *mansoni-*infected *B. pfeifferi* [23]. Fibrinogen-related proteins (FREPs) 1 and 2 were both up-regulated after niclosamide exposure; FREP2 was also up-regulated in S. mansoni-shedding *B. pfeifferi* [23]. Dermatopontin, a parasite-responsive gene frequently noted in studies of both *B. glabrata* and *B. pfeifferi*, was also up-regulated following niclosamide exposure.

A conspicuous response was the high up-regulation of over 100 diverse transcripts identified as LBP/BPI1 (lipopolysaccharide binding protein/bacterial permeability-increasing protein 1) in *B. pfeifferi* after exposure to niclosamide. LBP/BPI1 is an antimicrobial molecule found in the albumen gland of *B. glabrata* and egg masses [59]. Silencing of LBP/BPI1 expression in *B. glabrata* resulted in significant reduction of egg-laying, and death of eggs attributable to oomycete infections, providing evidence that LBP/BPI is involved in parental immune protection of offspring [60].

Transcripts homologous to *B. glabrata* tyrosinases (Tyr) 1, 2, and 3, are also up-regulated in response to niclosamide. In early-stage pre-patent *S. mansoni* infections Tyr-1 is up-regulated, and Tyr-3 is down-regulated in *B. pfeifferi* harboring cercariae-producing sporocysts [24]. Tyrosinases are involved in melanin synthesis and additionally might mark an early phase in initiation of castration by diverting tyrosine towards the production of melanin instead of dopamine in S. mansoni-infected *B. pfeifferi* [23]. Like LBP/BPI1, tyrosinase has also been isolated from *B. glabrata* egg masses and is presumed to provide an immunoprotective effect for developing embryos by contributing to the melanization of the egg membrane [59,61]. The additional considerable effort by the snail to make two egg mass-associated proteins in response to niclosamide is baffling, but might represent a last-ditch attempt to produce offspring before death. Alternatively, perhaps this is best viewed as an example of relatively non-specific innate immune responses that can be invoked by exposure to an unusual stressor, even if it is of an abiotic nature. Another consideration is that it represents a response to the presence of bacteria in the snail that might appear due to impaired hemocyte function or possibly due to failure to contain the gut microbiome in its usual compartment.

Another unexpected response was the high up-regulation of myomodulin-like neuropeptide in niclosamide-treated *B. pfeifferi.* Myomodulins are neurotransmitters involved in regulating feeding behavior by controlling radula protractor muscles used for feeding [62] in *Lymnaea stagnalis* [63] and *Aplysia californica* [64]. Myomodulin is down-regulated in prepatent *S. mansoni*-infected *B. glabrata* and this was implicated as possibly diminishing feeding efficiency in infected snails [65]. Down-regulation of a *B. pfeifferi* feeding circuitry peptide was seen in early and patent *S. mansoni* infections [23]. The up-regulated myomodulin activity noted provides evidence that basic physiological activities such as feeding are altered after niclosamide exposure. The mussel *Mytilus edulis* shows a decreased rate of feeding after exposure to hydrophobic organic chemicals, organochlorine compounds, organophosphate and carbamate pesticides, and pyrethroids [66–67].

With respect to features down-regulated following niclosamide exposure, it would seem transcription and translation efficiency would be hindered as evidenced by down-regulation of nearly a dozen ribosomal proteins, transcription factors, and mitogen-activated protein kinases (MAPKs). Of transcripts associated with stress responses, HSP 30 and HSP 70 cytosolic isoform were down-regulated along with an HSP 12 isoform. Neuroglobins are members of the hemoglobin superfamily of oxygen carriers, are expressed in the glial cells surrounding neurons and have been found in marine, freshwater, and terrestrial molluscs including the gastropods *Lymnaea stagnalis, Planorbis corneus, Aplysia californica, Helix pomatia* and *Cepaea nemoralis* [68]. Although we did not observe down-regulation of the hemoglobin-encoding gene noted by Zhang et al. [16] following exposure of *B. glabrata* to niclosamide, down-regulation of neuroglobin in niclosamide-exposed *B. pfeifferi* was observed. This could be associated with reduced availability of oxygen, at least for neural cells.

Significant down-regulation of a Cu-Zn SOD (−9.3 log_2_FC) in *B. pfeifferi* indicates that SODs have a more complex response to niclosamide than previously thought from the microarray study by Zhang et al. [16]. High expression of certain alleles of Cu-Zn SOD have been implicated in resistance of *B. glabrata* strain 13-16-R1 to *S. mansoni* [69–71] so it is not unlikely that different Cu-Zn SODs show distinctive responses to other stressors like niclosamide. Calmodulins, ubiquitous calcium-dependent signaling proteins responsible for regulating the uptake, transport, and secretion of calcium in gastropod shell formation [72–73], are expressed by *B. glabrata* in response to gram (-) and gram (+) bacteria, yeast [74], and in *B. glabrata* snail plasma containing larval *S. mansoni*. Here, we saw down-regulation of calmodulin in *B. pfeifferi* exposed to the niclosamide, raising the possibility that calmodulin expression is more responsive to biotic challenges. Transcripts related to cell adhesion like spondins that are expressed in *Biomphalaria* hemocytes [75] are also down-regulated.

### Responses of *B. pfeifferi* with cercariae-producing *S. mansoni* infections to sublethal niclosamide treatment

As previously noted, snails exposed to the combined effects of the biological stressor *S. mansoni* and the abiotic stressor niclosamide were surprisingly responsive (Fig 1), exhibiting large numbers of uniquely up- and down-regulated features, with many of these only modest in the degree of their differential expression. Among the more notable responses were several features associated with managing cell death in damaged tissues (Table 3). The transmembrane transporter ABCA3 is associated with resistance to xenobiotics and engulfment during apoptosis [76]. The enzymes glutaredoxin-2-like and catalase-like are both involved in reduction of hydrogen peroxide that may be released during niclosamide-induced apoptosis. An increase in apoptosis could account for the up-regulation of lysosomal endopeptidases such as cathepsin-L-like. Two mitochondria-associated transcripts that also play a role in gluconeogenesis, glyceraldehyde-3-phosphate dehydrogenase (GAPDH) and glycerol-3-phosphate dehydrogenase-like (GPDH) were also up-regulated. GAPDH accumulates in mitochondria during apoptosis and induces pro-apoptotic mitochondrial membrane permeability [77]. Niclosamide has been screened as a potential promoter of mitochondrial fragmentation by disrupting membrane potential, reducing ATP levels, and inducing apoptosis by caspase-3-activation in HeLa cells [78].

**Table 3.** Transcripts up-regulated in response to dual stressors (*S. mansoni* infection and sublethal niclosamide exposure) identified for their potential role in responding to programmed cell death. Except where noted, functions were obtained from Entrez Gene at http://www.ncbi.nlm.nih.gov/gene and UniProtKB at www.uniprot.org/uniprot.

| Transcript Description | Function |
| --- | --- |
| ABCA3 transmembrane transporter | Resistance to xenobiotics and engulfment during apoptosis |
| Growth arrest-specific protein 2-like | Cell cycle arrest; regulation of cell shape; may act as a cell death substrate for caspases |
| Glutaredoxin-2-like | Mitochondrial; response to hydrogen peroxide and regulation of apoptosis caused by oxidative stress |
| Calmodulin 2/4-like, 5, A-like | Can mediate the stress response calcium-dependent signaling that controls a variety of enzymes, ion channels, proteins, kinases, and phosphatases |
| Heparanase-like | Facilitates cell migration associated with metastasis, wound healing and inflammation |
| Catalase-like | Reduction of hydrogen peroxide |
| Caspase 3 and 8-like | TNF binding; endopeptidase activity involved in apoptosis |
| Tumor necrosis factor (TNF) and receptor | Induces cell death |
| Cathepsin-L-like | Lysosomal endopeptidase |
| Glyceraldehyde-3-phosphate dehydrogenase (GAPDH) | Induces pro-apoptotic mitochondrial membrane permeability (Deniaud et al. 2007) |

Pattern recognition receptors (PRRs), key elements responsible for the recognition of pathogens, showed mixed responses. Four distinct PRR genes were up-regulated: peptidoglycan-recognition protein SC2-like, ficolin-like, FREP 2, and FREP 10. We have reported the up-regulation of FREP 2 in *S. mansoni*-infected *B. pfeifferi* (Buddenborg et al 2017) but here we see four additional isoforms of FREP 2 up-regulated. Toll-like receptors (TLRs) which are involved in recognizing pathogens and activating conserved innate immune signaling pathways [79], were conspicuously down-regulated (TLRs 3, 4, 5, 7, and 8). Additional transcripts that function in various aspects of innate immune responses and that were down-regulated are C3 PZP-like alpha-2-macroglobulin domain-containing protein 8, hemolymph trypsin inhibitor B-like, tyrosine-3-monooxygenase, DBH-like monooxygenase 2, and tyramine beta-hydroxylase-like.

As with snails exposed to niclosamide alone, once again a down-regulation of transcripts for ribosomal proteins was noted. Reduction in ribosome production can be considered a stress response because it is a rapid and effective response against misfolded proteins [80] but may simply be an indication of a downgrading of general condition. Other down-regulated transcripts show diverse functional activity. Several annexins, intracellular Ca^2+^ and phospholipid binding proteins are down-regulated showing the possible disruption of regulation of membrane organization, trafficking, and the regulation of Ca^2+^ concentrations within cells [81].

Unlike the general up-regulation of CYPs in *B. pfeifferi* exposed only to niclosamide, *B. pfeifferi* with dual stressors highly down-regulate several CYPs (microsomal CYPs 2J1-like, 2B4-like, 3A29-like, 26A1-like, and mitochondrial CYP12A2-like). Mitochondrial CYP12A2-like is known to metabolize a variety of insecticides and xenobiotics [82]. We cannot discount that contribution to the down-regulation of this particular CYP is a result of mitochondrial degradation caused, in part, by niclosamide as noted previously as well as the additional stress of a patent *S. mansoni* infection.

## CONCLUDING REMARKS

This study provides a distinctive and detailed view of the nature of the response of field-derived *B. pfeifferi* to relevant stressors likely to be encountered in its environments, including infections with *S. mansoni*, just one of several digenetic trematodes known to commonly infect this snail in Africa [83], and exposure to the commonly used molluscicide, niclosamide. It is important to gain additional detailed information regarding the effects of niclosamide on snails, particularly those that harbor schistosome infections. For example, do infected snails succumb more readily to treatment and if so, why? This particular aspect of molluscicide use has not been widely investigated.

In general, exposure to niclosamide alone resulted in the fewest responsive features in *B. pfeifferi* (1,711) followed by infection with *S. mansoni* (2,271) and then by the combination of niclosamide and *S. mansoni* (7,683). Snails in these three groups all responded in very distinct ways, but in each case with more features up- than down-regulated. Sublethal exposure to a single xenobiotic provoked about 67% as large a transcriptomic response as was noted for snails shedding *S. mansoni* cercariae, snails that had probably been infected with the parasite for at least a month and harbored large numbers of daughter sporocysts. The fact that snails that received the combination of infection and niclosamide responded so much more vigorously with so many distinctive features suggests that they were under greater duress and that their responses in some sense preempted the responses of snails in the other two groups.

Exposure to niclosamide alone provoked up-regulation of several features associated with response to xenobiotics including cytochrome p450s, heat shock proteins, multidrug resistant transporters and glutathione-S-transferases, confirming many of the observations made by Zhang et al [16] in a microarray study of *B. glabrata* exposed to sublethal doses of niclosamide. Several additional unique aspects of the response to niclosamide were also noted given the increased resolution provided by Illumina sequencing. We note that one of the effects of niclosamide on *B. pfeifferi* may be to contribute to redox imbalance because glutathione is being used by glutathione-S-transferases to conjugate xenobiotics but may not be sufficiently regenerated because of down-regulated activity of glutathione reductase.

Exposure of infected snails to niclosamide was noteworthy in revealing the involvement of several features not found to be responsive to either stressor alone. Although many of the uniquely expressed features did not respond dramatically, the ones that did were indicative of responses associated with apoptosis, reduced protein synthesis, reduced production of some CYPs and thus diminished detoxification ability, and diminished innate immune function. Accordingly, we hypothesize that the combination of stressors was likely overcoming the snail’s ability to maintain homeostasis. The snail mounts a considerable transcriptomic response to the presence of cercariae-producing sporocysts [23] and it is not hard to imagine that the energy demand placed on infected snails by continual production of cercariae takes an additional toll. The mortality rate of *B. pfeifferi* infected with *S. mansoni* is significantly higher than that noted for unexposed control snails [25]. The molluscicide-exposed infected snails selected for sequencing were alive when sampled, but the transcriptional profiles suggested they were not thriving. This is broadly in agreement with observations made to indicate that *B. sudanica* with *S. mansoni* infections succumb to sublethal niclosamide treatment at a higher rate than do uninfected controls [20]. In other words, the combination of stressors used here exposed the limits of what these snails can do to maintain homeostasis.

Even though *S. mansoni* sporocysts within snails exposed to niclosamide expressed more transcripts than in untreated snails, there was little about the response to suggest they possessed any distinctive or large-scale ability to respond to a xenobiotic like niclosamide. Furthermore, the sporocyst response did not appear to be as indicative of a failure to maintain homeostasis as we noted for snails. This is in keeping with the general observation that the lethal dose of niclosamide for sporocysts is probably much higher than for snails [20]. Although it is clear that both miracidia and cercariae are vulnerable to niclosamide [17–18], this may be a reflection of their more aerobic metabolism and that they would be more fully exposed to the action of niclosamide as compared to sporocysts nested within the tissues of an infected snail. Although we did not observe a strong negative effect of molluscicide exposure on the transcriptomics responses of sporocysts, given the relatively unhealthy state of the snail that we detected, it would inevitably follow that the condition of the sporocysts would degenerate.

In conclusion, we noted remarkably distinctive transcriptomics responses for *B. pfeifferi* depending on the nature of the stressor treatment they received, and that the combination of niclosamide and *S. mansoni* infection imposed a level of stress on the snails that resulted in a massive response comprised of many features we had not observed previously. This study contributes to the growing list of molecular participants that may govern the outcomes of the intimate interrelationships between snails and schistosomes, and that may help us understand how snail host biology might be targeted for disruption by molluscicidal chemicals.

